# Dissociable Roles of Primary Motor and Supplementary Motor Cortex in Shaping the Neural Drive to Muscle

**DOI:** 10.1101/2025.10.28.685170

**Authors:** Yori R. Escalante, Yuming Lei

**Author notes:** Correspondence to, Yuming Lei, PhD, Department of Kinesiology & Sport Management, Texas A&M University.

## Abstract

The primary motor cortex (M1) and supplementary motor area (SMA) are critical for motor execution and planning, yet their distinct causal contributions to modulating the neural drive to muscles remain incompletely understood. To dissociate their roles, we applied inhibitory transcranial magnetic stimulation (TMS) over M1, SMA, or a sham condition in 72 healthy participants and characterized the activity of single motor units from high-density EMG recorded during a sustained isometric contraction. Our results revealed a clear functional divergence. M1 inhibition produced a direct failure of motor output, causing a rapid force decline compared to sham, which was strongly correlated with a reduction in motor unit firing rates. Conversely, SMA inhibition did not impair net force. Instead, it altered the fundamental structure of the motor command, compelling a compensatory strategy characterized by a reliance on smaller-amplitude motor units with lower firing rates and a marked degradation of the low-frequency (delta-band) coherence that organizes stable output. These results provide direct causal evidence that M1 directly dictates the magnitude of motor output via population firing rates, while SMA orchestrates the composition and temporal structure of the active motor unit pool to generate an efficient and stable command.

**Significance Statement:** Using causal brain stimulation, we provide the first direct evidence that the primary motor cortex (M1) and supplementary motor area (SMA) serve distinct, non-redundant roles in controlling motor unit populations. We show that inhibiting M1 directly impairs force magnitude by reducing motor unit firing rates. In contrast, inhibiting SMA spares net force but disrupts the underlying motor plan, compelling a compensatory strategy that uses smaller motor units and degrades temporal firing organization via a loss of delta-band coherence. By demonstrating that M1 governs motor power while SMA organizes motor strategy, this study opens new avenues for personalizing neuro-rehabilitation to address the specific cortical origin of a patient’s motor deficits.

## Introduction

The ability to generate and sustain precise muscular force is fundamental to motor control, enabling everything from grasping a delicate object to maintaining posture against gravity (Evarts, 1968; Wolpert & Ghahramani, 2000). This precise control is orchestrated by a complex cortical architecture that translates high-level goals into an organized neural drive to muscles (Houk & Wise, 1995; Farina et al., 2014; Battaglia-Mayer & Caminiti, 2019). Decades of research have established a hierarchical model in which the primary motor cortex (M1) and the supplementary motor area (SMA) serve as two critical, yet functionally distinct, nodes (Penfield & Rasmussen, 1950; Nachev et al., 2008; Omrani et al., 2017). M1 is widely regarded as the principal output node of the motor cortex, executing voluntary movements through its dense corticospinal projections (Lemon, 2008; Omrani et al., 2017). In contrast, the SMA is considered a higher-order area, essential for the planning, sequencing, and temporal organization of movements, particularly those that are internally generated rather than guided by external cues (Roland et al., 1980; Tanji, 2001).

While this functional distinction is well-supported by a wealth of correlational evidence from neuroimaging and electrophysiology, it harbors a fundamental ambiguity (Nachev et al., 2008; Omrani et al., 2017). Correlational methods cannot definitively establish the unique causal contributions of each region to the final motor command (Hallett, 2007). Crucially, it remains unknown how the abstract roles of "execution" and "planning" are mechanistically implemented at the level of the spinal motor neuron pool, the final common pathway for motor control (Sherrington, 1906; Heckman & Enoka, 2012). The neural drive to a muscle is ultimately translated into force through two fundamental parameters— which motor units are recruited and their respective firing rates (Henneman et al., 1965; Enoka & Fuglevand, 2001). Therefore, a comprehensive understanding of cortical motor control requires bridging this explanatory gap, moving beyond high-level descriptions to a causal, mechanistic account of how M1 and SMA distinctly shape the activity of the active motor unit pool to generate a stable and effective motor command.

To bridge this gap, a causal, "perturb-and-measure" approach is required (Paus, 2005). Transcranial magnetic stimulation (TMS) offers a powerful tool to achieve this, allowing for the creation of a temporary, reversible "virtual lesion" over a targeted cortical area, thereby establishing the causal necessity of that region for a given function (Hallett, 2000; Walsh & Cowey, 2000). In parallel, advances in high-density surface electromyography (HD-EMG) and blind-source separation algorithms now permit the non-invasive decomposition of the interferential surface signal into the constituent action potential trains of dozens of single motor units (Holobar & Zazula, 2007; Negro et al., 2016). Combining these powerful techniques offers a unique opportunity to causally dissect the cortical motor command, enabling us to ask a precise question: if M1 and SMA are functionally dissociated, how does inhibiting each region differentially alter the composition, firing rate dynamics, and temporal coherence of the active motor unit pool during a sustained contraction?

This study was designed to test two distinct hypotheses, each grounded in the canonical roles of M1 and SMA. First, we hypothesized that inhibiting M1, as the principal execution node, would result in a direct failure of motor output. We predicted this impairment would manifest as a decline in force directly attributable to a reduction in motor unit firing rates. Conversely, we hypothesized that inhibiting SMA, as a higher-order planning and organizational hub, would spare the overall magnitude of force but disrupt its underlying structure and efficiency. We predicted this disruption would compel a compensatory neural strategy, characterized by a reorganization of the recruited motor unit pool and a degradation of the low-frequency coherence that is thought to organize a stable neural drive. By testing these hypotheses, this study provides direct causal evidence for a functional dissociation in how these two critical motor areas shape the final neural drive to muscle.

## Methods

### Participants

A total of 72 healthy, right-handed individuals (age range: 18–30 years) were recruited from the university and local community. All potential participants underwent a thorough screening process to ensure they met the study’s inclusion criteria. Exclusion criteria were designed to minimize confounding variables and ensure safety. These included: (1) any self-reported history of neurological disorders such as stroke, Parkinson’s disease, epilepsy, or peripheral neuropathy; (2) any current or recent musculoskeletal injuries, pain, or conditions affecting the right arm, hand, shoulder, or neck that could interfere with task performance; (3) the use of any psychoactive medications or drugs known to alter cortical excitability or motor control; and (4) any contraindications to Transcranial Magnetic Stimulation (TMS) as per established safety guidelines (e.g., presence of metal implants in the head, personal or family history of seizures). The experimental protocol was formally reviewed and approved by the Institutional Review Board (IRB) of Texas A&M University. The study was conducted in full compliance with the ethical principles outlined in the Declaration of Helsinki. Prior to any experimental procedures, all participants were provided with a detailed explanation of the study’s purpose, procedures, potential risks, and benefits, after which they provided written informed consent.

### EMG recording

To characterize the activity of individual motor units, high-density surface electromyography (HDsEMG) was used to record myoelectric signals from the right First Dorsal Interosseous (FDI) muscle. To ensure a high signal-to-noise ratio, the skin overlying the muscle belly was carefully prepared. This procedure involved gentle abrasion with a preparatory gel followed by cleansing with an alcohol solution to minimize skin impedance. The Delsys Trigno Galileo sensor (Delsys, Natick, MA, USA), featuring a 4-electrode array, was then securely affixed over the prepared area, aligned with the presumed orientation of the muscle fibers. The raw HDsEMG signals were sampled at a rate of 2222 Hz and wirelessly transmitted to the acquisition software. This high-density recording configuration was specifically chosen to facilitate the subsequent offline decomposition of the interferential surface signal into the constituent action potential trains of single motor units. For the purpose of TMS hotspotting and the determination of the resting motor threshold (RMT), a separate, standard EMG setup was used. Disposable bipolar Ag-Cl surface electrodes were placed over the FDI muscle in a belly-tendon montage. The active electrode was positioned over the muscle belly, while the reference electrode was placed on the metacarpophalangeal joint of the index finger. A ground electrode was secured on a bony prominence of the right wrist (the ulnar styloid process). This configuration is optimal for recording the clear, large-amplitude motor-evoked potentials (MEPs) required for precise TMS localization.

### Transcranial magnetic stimulation (TMS) protocols

A Magstim Rapid^2^ stimulator connected to a 70mm figure-of-eight coil (Magstim, Whitland, UK) was used for all stimulation procedures. To ensure spatial accuracy and consistency across all procedures, coil positioning was guided and monitored in real-time using a Brainsight Neuronavigation system (Rogue Research, Montréal, QC, Canada). The cortical representation of the FDI muscle was precisely located prior to the main experiment. With the coil held tangentially to the scalp at a 45-degree angle to the mid-sagittal line (inducing a posterior-anterior current), single TMS pulses were delivered over the contralateral primary motor cortex. The "motor hotspot" was identified as the scalp location that consistently produced the largest MEPs in the resting FDI muscle. This location was marked within the neuronavigation system and used as the target for all subsequent M1 stimulation. We used an inhibitory, low-frequency (1-Hz) repetitive TMS (rTMS) protocol to modulate cortical excitability. To ensure the stimulation was subthreshold, the intensity was set to 40% of each individual’s Resting Motor Threshold (RMT). The RMT was determined beforehand at the motor hotspot and defined as the minimum intensity required to elicit an MEP of at least 0.05 mV peak-to-peak amplitude in at least 5 of 10 consecutive trials. This low intensity prevents direct muscle twitches that could otherwise contaminate the EMG signal or interfere with the voluntary motor command. Participants underwent one of three rTMS conditions in a between-subjects design. All stimulation sites were precisely localized and monitored using a Brainsight Neuronavigation system. (1) Primary Motor Cortex (M1) Stimulation: The coil was precisely maintained over the pre-determined FDI motor hotspot. This condition was designed to directly probe the role of the primary corticospinal output pathway in sustaining the motor command. (2) Supplementary Motor Area (SMA) Stimulation: The coil was positioned 4cm anterior to the vertex (Cz) along the sagittal midline, a standard landmark-based approximation for the human SMA (Arai et al., 2012). This condition was designed to investigate the SMA’s role in higher-order motor functions, such as motor planning and reliance on an internal motor representation when visual feedback is absent. Crucially, this stimulation site is functionally and spatially distinct from the M1 target. Recent work from our group has demonstrated the high spatial focality of stimulation at this location; a facilitatory connection between SMA and M1 was present only at the 4 cm site and was absent at sites just 1 cm more anterior (Kim et al., 2025). This spatial specificity provides strong evidence that the rTMS applied over the SMA in the current study modulated a distinct neural population from that targeted by direct M1 stimulation, minimizing the possibility of overlapping effects. (3) Sham Stimulation: A sham condition was used to account for auditory and somatosensory effects associated with TMS. The coil was placed over the vertex but angled perpendicularly with only its edge touching the scalp. This orientation replicates the clicking sound and scalp sensation of real TMS without inducing a significant electrical field in the underlying cortical tissue.

### Experimental Paradigm

Participants were comfortably seated in an adjustable chair with their right forearm securely resting on a padded surface. This setup was designed to stabilize the limb and isolate the actions of the hand and wrist. Force output from the FDI muscle, generated via index finger abduction, was measured using a Futek Load Cell (Futek, Irvine, CA, USA). The analog force signal was digitized and recorded by a custom MATLAB script (Mathworks, Natick, MA, USA), which also controlled the real-time visual display. Prior to the main task, each participant’s maximal voluntary contraction (MVC) was established to normalize task difficulty. Participants were instructed to perform three maximal isometric contractions, each lasting 3-5 seconds, with a 60-second rest period between efforts to prevent fatigue. The highest absolute force value achieved across these three trials was recorded as that individual’s MVC. The main task required participants to produce and maintain a steady, submaximal force at 40% of their individual MVC. Each 50-second trial was precisely structured into two sequential phases: (1) Visual Feedback (FB) Phase (0–20 seconds): This initial phase consisted of a 20-second period where participants had full visual feedback. The phase was further structured to include a 10-second baseline for the participant to orient to the cursor, followed by a 5-second ramp-up to the target, and a final 5-second stable hold. Throughout this phase, a real-time cursor representing the participant’s force output was displayed, and they were instructed to keep it precisely aligned with a target line representing 40% MVC; (2) No Visual Feedback (NFB) Phase (20–50 seconds): Immediately upon cessation of visual feedback, the NFB phase began. The cursor was removed from the screen, and simultaneously, 1-Hz rTMS was initiated and delivered for the entire 30-second duration (Figure 1A). Participants were explicitly instructed to maintain the same level of force for the remaining 30 seconds. This study employed a between-subjects design where participants were randomly assigned to one of three experimental groups. Each participant performed two practice trials to familiarize themselves with the task dynamics before proceeding to three recorded test trials under their assigned stimulation condition (M1, SMA, or Sham; Figure 1B). Critically, to mitigate the effect of trial-to-trial learning, no performance feedback was provided between the recorded trials.

**Figure 1.**
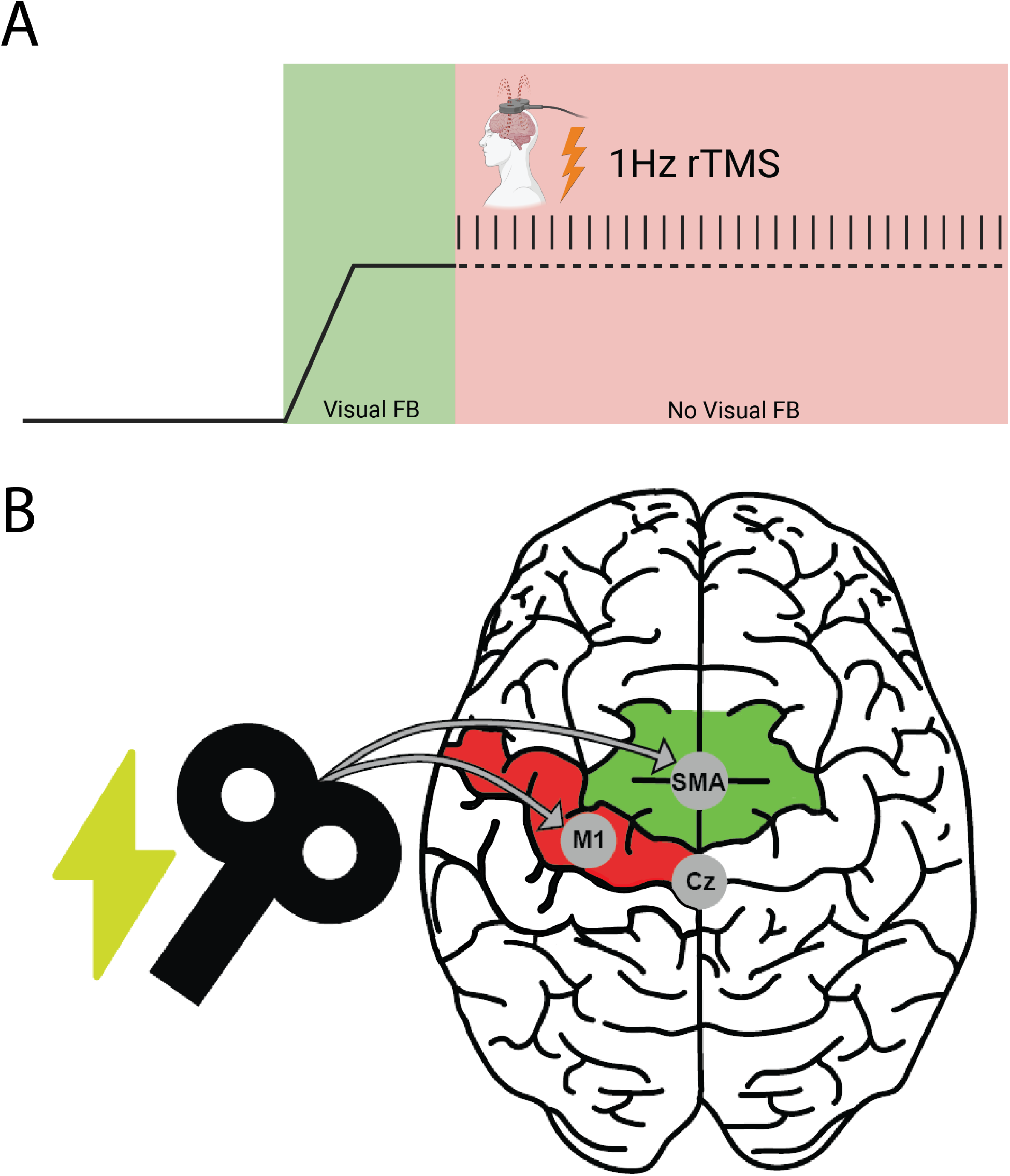
Experimental Paradigm and Stimulation Sites. This figure provides a schematic overview of the experimental task and the targeted brain regions. (**A**) The temporal structure of a single trial. Participants first performed a force-matching task with continuous visual feedback (FB, green area), which included a ramp and hold phase. Subsequently, the visual feedback was removed (NFB, red area), and participants were instructed to maintain the same force level. Repetitive transcranial magnetic stimulation (1-Hz rTMS) was delivered for the entire 30-second duration of the NFB phase. (**B**) Illustration of the transcranial magnetic stimulation (TMS) setup. The figure-of-eight coil was positioned over the primary motor cortex (M1, red area) or the supplementary motor area (SMA, green area) to causally probe their roles in motor control. The vertex (Cz) is shown as a key landmark for localizing the SMA.

### Force Data Processing

The raw analog force signal, recorded in millivolts from the load cell, was digitized for offline analysis. To ensure comparability across participants, this signal was then normalized and expressed as a percentage of each individual’s MVC (%MVC), resulting in a continuous time series of force output for each trial. From this time series, we calculated two primary dependent variables: (1) Mean Force (%MVC): The average force maintained during a given period; (2) Force Variability (%MVC-CoV): The stability of the force output, quantified using the coefficient of variation. These dependent variables were compared both within each group (between the FB and NFB phases) and between the three experimental groups (M1, SMA, and Sham).

### Motor Unit Data Processing

Single motor unit action potential trains were identified from the raw HDsEMG signals using the validated Delsys Neuromap decomposition algorithm (Hu et al., 2012; Lei et al., 2018). To ensure a standardized analysis window across all trials and participants, the resulting spike trains for each trial were manually trimmed to the required 50-second period. This was guided by the consistent activity of the first-identified motor unit, which was active throughout the entire task duration for all participants, thereby providing a reliable temporal anchor. From these trimmed spike trains, the instantaneous firing rate of each motor unit was calculated as the inverse of its inter-spike intervals (ISIs). From these processed spike trains, three primary motor unit metrics were calculated for each participant, analogous to the force data analysis: (1) Mean Motor Unit Firing Rate (MUFR): To derive a single measure of neural drive, a multi-step averaging process was used. First, the instantaneous firing rates of all concurrently active motor units were averaged to create a single continuous MUFR time series for each trial. This time series was then averaged across the FB and NFB phases, resulting in a single mean MUFR value for each phase per participant. (2) MUFR Variability (MUFR-CoV): The stability of the neural drive was quantified by calculating the coefficient of variation of the MUFR time series for both the FB and NFB phases. (3) Motor Unit Action Potential (MUAP) Amplitude: The peak-to-peak amplitude of each motor unit’s waveform (its spike-triggered average) was provided by the NeuroMap decomposition software (Escalante et al., 2025). These individual amplitudes were then averaged across all identified motor units within a trial, and finally averaged across the three test trials, resulting in a single mean MUAP amplitude value for each participant.

### Motor Unit Coherence Analysis

To assess the temporal organization of the neural drive and quantify the strength of common synaptic input to the motoneuron pool, we performed a motor unit coherence analysis. This frequency-domain measure quantifies the degree of correlated firing among different motor units, which directly reflects shared, oscillatory input from the central nervous system. The analysis was performed on the motor unit spike trains identified during the 30-second NFB phase. First, to convert the discrete spike train data into a continuous signal suitable for spectral analysis, each spike train was smoothed by convolution with a 400-ms Hanning window. Next, the pairwise magnitude-squared coherence between all combinations of these smoothed spike trains was computed for each participant using the mscohere function in MATLAB (Bao et al., 2022; Bao & Lei, 2024). This function estimates the correlation between the signals at specific frequencies. To ensure the coherence values were normally distributed for valid statistical comparison, the resulting values (C) were transformed using Fisher’s z-transformation with the formula:

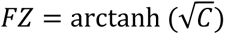

The transformed coherence values (FZ) for all motor unit pairs were then averaged for each participant. Finally, these mean coherence values were calculated within three canonical frequency bands of interest, each chosen for its distinct and well-established neurophysiological origin. Coherence in the delta band (0.5–5 Hz) is primarily thought to reflect the low-frequency "common drive" to the motoneuron pool, which is essential for stabilizing force output during steady contractions (De Luca et al., 1982). The alpha band (5–13 Hz), in contrast, is more closely associated with physiological tremor and contributions from subcortical pathways (Elble & Koller, 1990). Lastly, coherence in the beta band (13–30 Hz) is strongly linked to descending corticospinal drive and is considered a key signature of oscillatory communication from the sensorimotor cortex (Conway et al., 1995).

### Statistical Analysis

All statistical analyses were performed using custom MATLAB scripts and SPSS (Version 28.0, IBM), with an alpha level for all tests set to p < 0.05. Prior to parametric testing, data were assessed for assumptions using the Shapiro-Wilk test for normality and Levene’s test for homogeneity of variances. First, to assess the effect of removing visual feedback within each experimental group (M1, SMA, Sham), two-tailed paired-samples t-tests were used to compare the mean values of %MVC, %MVC-CoV, MUFR, and MUFR-CoV between the FB and NFB phases. Next, to evaluate the dynamic effects of rTMS over time, the 30-second NFB phase was binned into six consecutive 5-second intervals. A two-way mixed-model repeated measures ANOVA was then performed on the key dependent variables (%MVC and MUFR), with Time (6 levels) serving as the within-subjects factor and Group (M1, SMA, Sham) as the between-subjects factor. For this ANOVA, Mauchly’s test was used to assess the sphericity assumption; if the assumption was violated, a Greenhouse-Geisser correction was applied. If a significant Time x Group interaction was revealed, we followed up with separate one-way ANOVAs at each time point, with any significant effects further interrogated using post-hoc tests with a

Bonferroni correction for multiple comparisons. One-way ANOVAs with Bonferroni-corrected post-hoc tests were also used to assess overall between-group differences in MUAP amplitude and motor unit coherence within each frequency band. Finally, to quantify the relationship between neural drive and force output, Pearson correlation coefficients (r) were computed for each experimental group, correlating the mean MUFR with the mean %MVC during the NFB phase.

## Results

### Baseline Characteristics

Prior to analyzing the primary experimental data, we performed a series of baseline comparisons to ensure that the randomly assigned experimental groups were equivalent on key demographic and neurophysiological measures. This step is critical to confirm that any observed differences in task performance can be confidently attributed to the effects of the rTMS intervention, rather than pre-existing physiological disparities. First, we assessed baseline muscle strength. A one-way ANOVA was conducted to compare the Maximal Voluntary Contraction (MVC) across the M1, SMA, and Sham groups. The results indicated that the mean MVC strength did not significantly differ between the M1 (Mean = 17.94 mV, Standard Deviation (SD) = 5.52), SMA (Mean = 16.47 mV, SD = 5.08), and Sham (Mean = 19.88 mV, SD = 5.99) groups, *F*(2,72) = 2.383, *p* = 0.1. This confirms that all three groups possessed comparable physical strength at the outset. Next, we evaluated baseline corticospinal excitability for the two groups receiving active stimulation. An independent samples t-test revealed no significant difference in the RMT between the M1 (Mean = 47.74% of Maximum Stimulator Output (MSO), SD = 8.79) and SMA (Mean = 51.65% MSO, SD = 10.53) groups, *t*(49) = −1.327, *p* = 0.2. This result indicates that the M1 and SMA groups began the experiment with similar baseline levels of cortical excitability. Taken together, these baseline analyses establish that the experimental groups were well-matched on both peripheral strength and central excitability, providing a solid foundation for attributing any subsequent differences in force control or motor unit activity to the specific effects of the rTMS protocols.

### Force Control is Impaired in the Absence of Visual Feedback

We first sought to confirm that the removal of visual feedback created a more challenging motor control state, thereby compelling participants to rely on internal control mechanisms. To achieve this, paired-samples t-tests were conducted within each experimental group to compare performance during the Visual Feedback (FB) phase to the No Visual Feedback (NFB) phase. The results, depicted in Figure 2, demonstrate a universal and significant degradation of motor performance when visual guidance was withdrawn. A significant decline in mean force was observed across all three experimental groups during the NFB phase. As shown in Figure 2A, this decline was statistically robust for the M1 group (*t*(24) = −5.196, *p* < 0.001, Cohen’s d = 1.039), the SMA group (*t*(24) = 3.824, *p* < 0.001, Cohen’s d = 0.765), and the Sham group (*t*(24) = 2.089, *p* = 0.02). This finding confirms a universal tendency for force output to drift downwards when participants must rely on internal representations of effort rather than external visual guidance. In parallel with the decline in magnitude, the stability of the force output also degraded significantly. This loss of stability was reflected by a marked and highly significant increase in the coefficient of variation of force for all three groups: M1 (*t*(24) = −9.099, *p* < 0.001), SMA (*t*(24) = −7.177, *p* < 0.001), and Sham (*t*(24) = −10.98, *p* < 0.001) (Figure 2B). Taken together, these results confirm that the No Visual Feedback condition successfully induced a more challenging motor state, characterized by both a reduced and a more variable force output.

**Figure 2.**
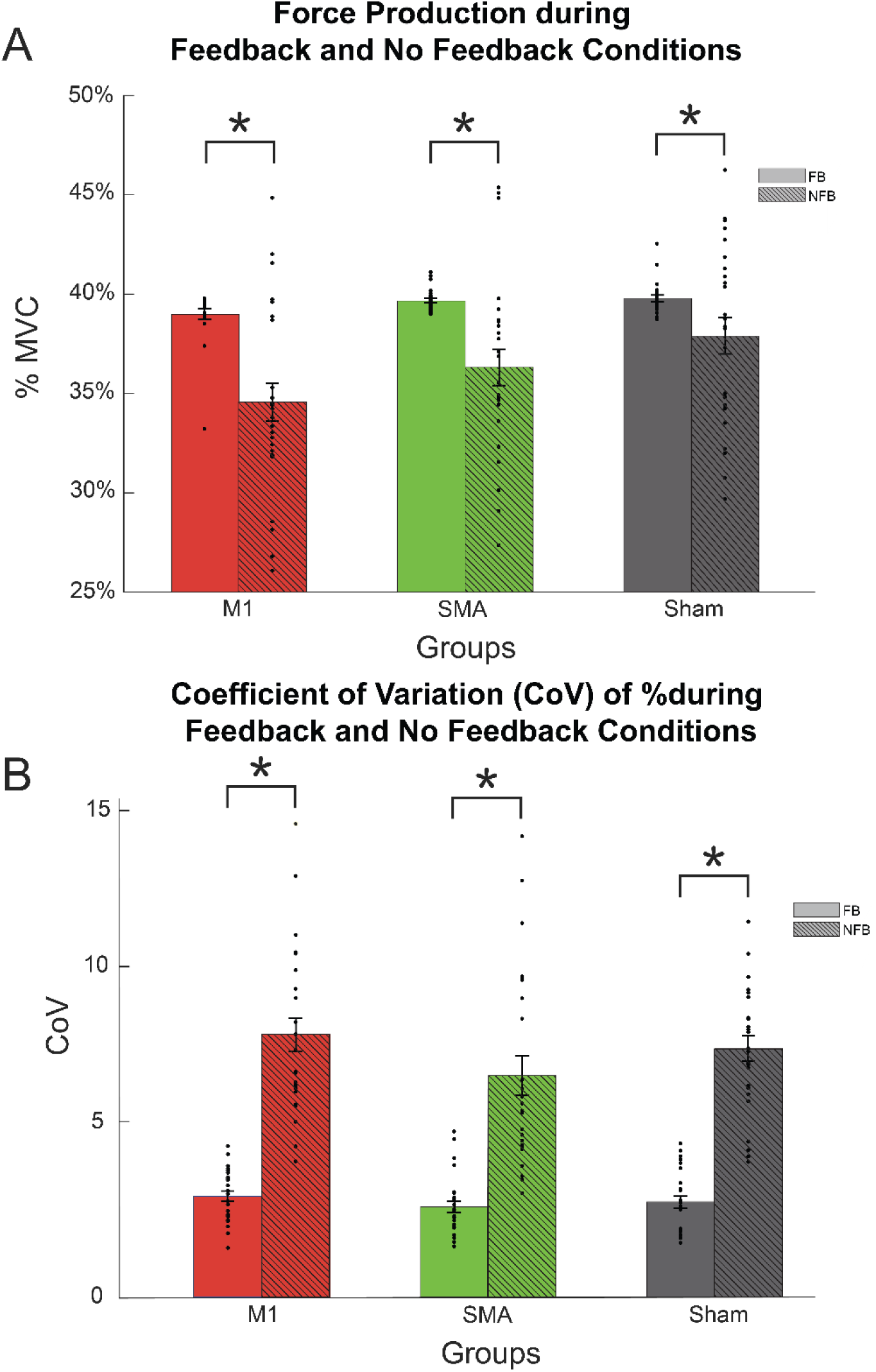
Removal of Visual Feedback Degrades Force Control. This figure illustrates the general behavioral effect of removing visual feedback, demonstrating that the task became more difficult for all participant groups. Data are shown as mean ± standard error of the mean (SEM), with individual participant data overlaid as dots. (**A**) Mean force production, expressed as a percentage of maximal voluntary contraction (%MVC), during the visual feedback (FB) and no visual feedback (NFB) phases. In all three groups, mean force was significantly lower during the NFB phase. (**B**) Mean force variability, quantified by the coefficient of variation (CoV), during the FB and NFB phases. In all three groups, force output was significantly more variable during the NFB phase. Asterisks (*) indicate a statistically significant difference (*p* < 0.05) between the FB and NFB phases within each group.

### M1 Inhibition, but Not SMA, Impairs Sustained Force Control

To isolate the specific, causal effects of each rTMS protocol on sustained force control, we analyzed the time course of force production (%MVC) during the NFB phase. The primary statistical test was a two-way repeated-measures ANOVA, with Time (six 5-second bins) as the within-subjects factor and Group (M1, SMA, Sham) as the between-subjects factor. The ANOVA, with a Greenhouse-Geisser correction applied due to a violation of sphericity (*χ²*(14) = 365.674, *p* < 0.001), revealed a significant main effect of time, *F*(1.508, 72) = 26.83, *p* < 0.001. This confirms the general downward drift of force over time for all groups. However, the overall Time x Group interaction did not reach statistical significance (*p* = 0.297). To test our specific a priori hypotheses regarding the differential effects of M1 and SMA stimulation relative to the Sham condition, we proceeded with planned one-way ANOVAs at each of the six 5-second time bins. As shown in Figure 3, these more sensitive analyses revealed a clear dissociation between the behavioral consequences of M1 and SMA stimulation. As hypothesized, inhibitory rTMS over M1 resulted in a significant impairment of force control compared to the Sham condition. The continuous force traces in Figure 3A illustrate a clear separation between the M1 and Sham groups. Post-hoc tests with Bonferroni correction on the binned data confirmed this observation, revealing that the M1 group produced significantly less force than the Sham group during the initial 20 seconds of the NFB phase. Specifically, these significant differences were observed at the 1-5s (*p* = 0.014), 5-10s (*p* = 0.007), 10-15s (*p* = 0.006), and 15-20s (*p* = 0.027) time bins (Figure 3B). This pattern indicates that M1 inhibition caused a rapid and robust decline in the ability to sustain the required force output.

**Figure 3.**
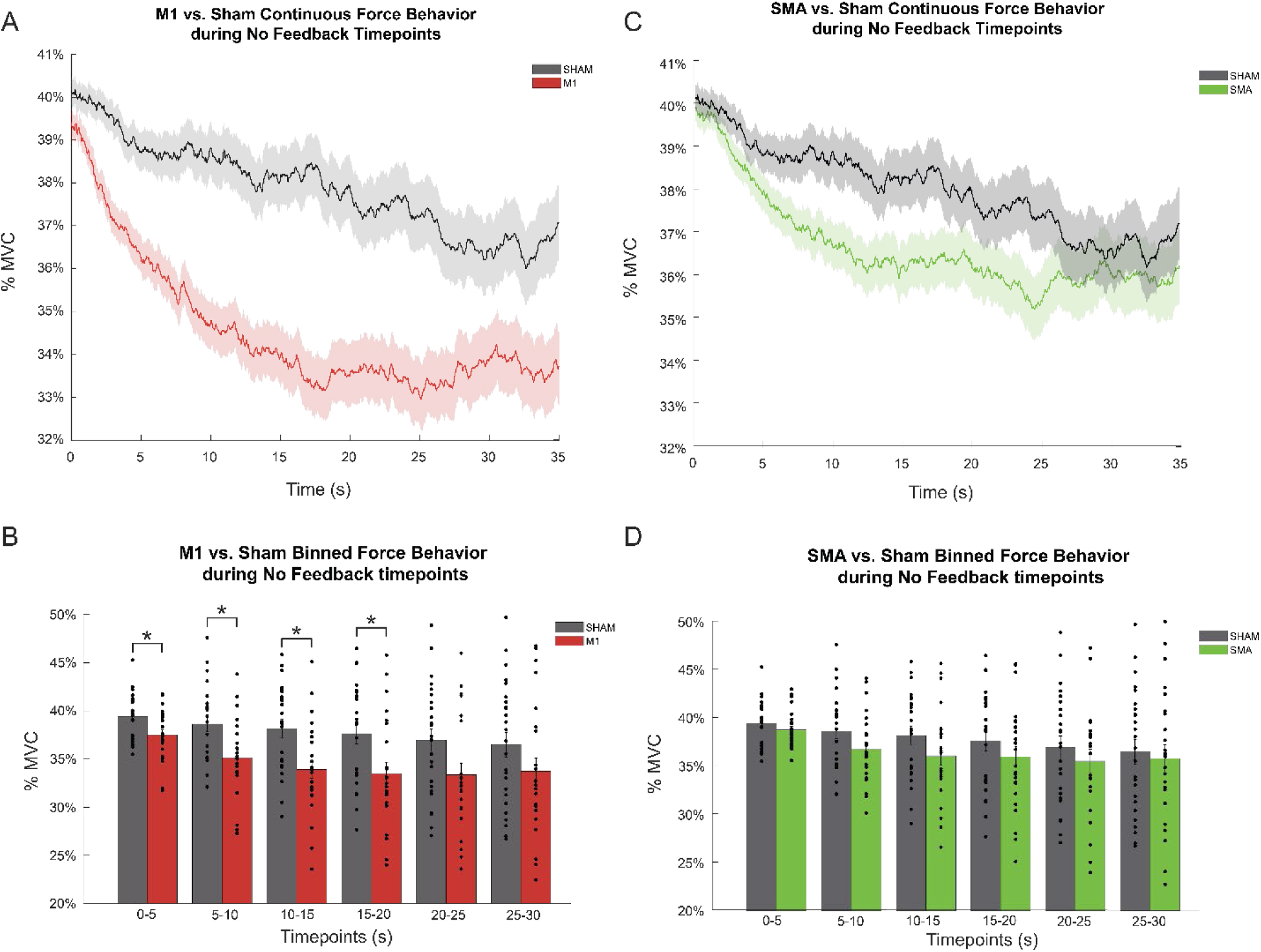
M1 Inhibition, but Not SMA, Causes a Significant Decline in Force. This figure presents the primary behavioral finding, dissociating the effects of M1 and SMA stimulation on sustained force control during the No Visual Feedback (NFB) phase. (**A**) The continuous time course of mean force (%MVC) for the M1 group (red) compared to the Sham group (black). Shaded areas represent SEM. (**B**) Binned force data for the M1 and Sham groups, averaged into 5-second intervals. Inhibition of M1 resulted in a significantly lower force output compared to Sham for the first 20 seconds of the NFB phase. (**C**) The continuous time course of mean force for the SMA group (green) compared to the Sham group (black). (**D**) Binned force data for the SMA and Sham groups. Inhibition of SMA did not result in a significant change in force magnitude compared to Sham at any time point. Error bars represent SEM, and asterisks (*) denote a significant difference (*p* < 0.05) between the stimulation group and the Sham group within that time bin.

In contrast to the effects of M1 stimulation, and consistent with our second hypothesis, rTMS over the SMA did not result in a significant loss of force magnitude (Figure 3C). The statistical analysis confirmed that there were no statistically significant differences in mean %MVC between the SMA and Sham groups at any of the six time bins analyzed (Figure 3D). This finding demonstrates that, unlike M1, the SMA is not essential for maintaining the overall magnitude of force, suggesting its role lies in a different aspect of motor control that is not captured by this behavioral measure alone.

### A Dissociation in Motor Unit Control Strategies

To understand the neural mechanisms underlying the observed behavioral dissociation, we analyzed the activity of 5,277 successfully decomposed motor units from the FDI muscle. First, we examined the general effects of removing visual feedback on motor unit activity. As shown in Figure 4A, a significant drop in MUFR from the FB to the NFB phase was observed in both the actively stimulated M1 (*p* < 0.001) and SMA (*p* < 0.001) groups, while the Sham group showed no significant change (*p* = 0.065). In addition to this specific effect on firing rate magnitude, a more general effect on firing rate stability was observed across all groups. Consistent with the behavioral data showing reduced force stability, the motor unit firing rate variability (MUFR-CoV) increased significantly from the FB to the NFB phase for all three groups (M1, SMA, and Sham, all *p* < 0.001) (Figure 4B).

**Figure 4.**
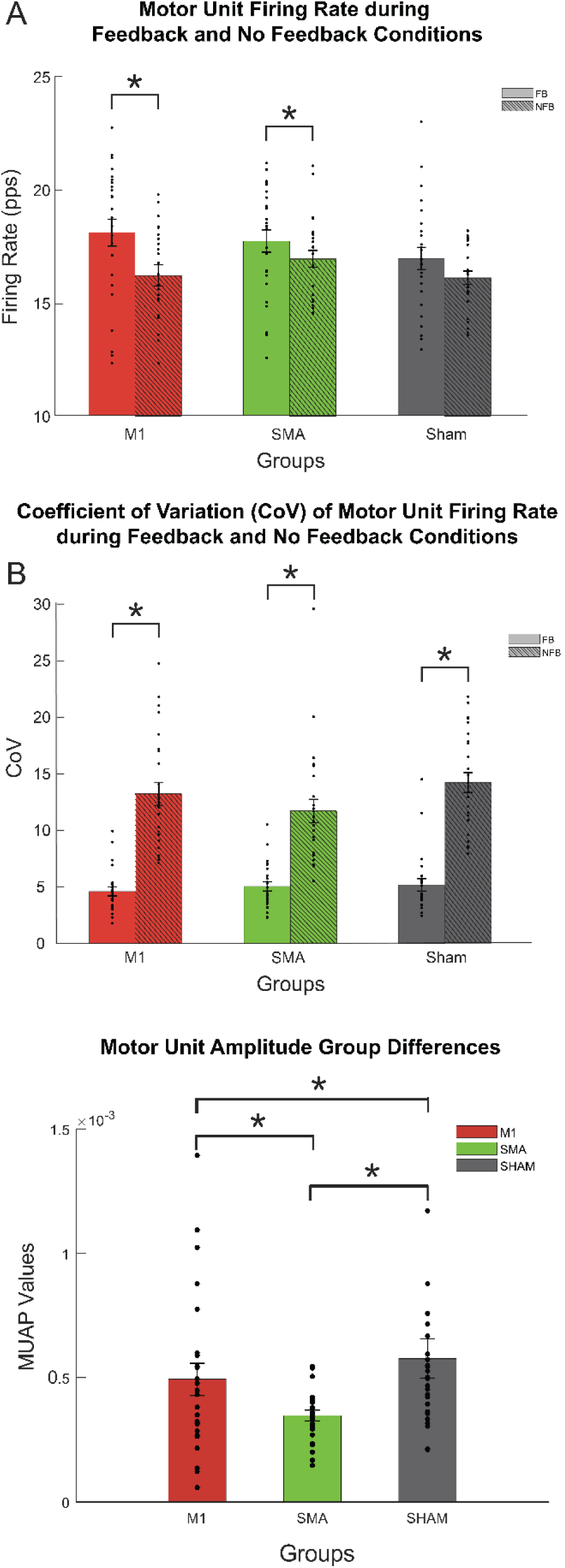
Dissociable Effects of rTMS on Motor Unit Firing and Recruitment. This figure details the neurophysiological effects of the interventions on three core motor unit properties. (**A**) Mean motor unit firing rate (MUFR) during the FB and NFB phases. Active stimulation over M1 and SMA, but not Sham, resulted in a significant decrease in MUFR. (**B**) Motor unit firing rate variability (MUFR-CoV). Firing rate became significantly more variable during the NFB phase for all three groups. (**C**) A between-group comparison of the average motor unit action potential (MUAP) amplitude, a proxy for motor unit size. All three groups were significantly different from one another, indicating that the rTMS protocols altered the composition of the recruited motor unit pool. Bars represent the group mean, error bars represent SEM, and asterisks (*) denote statistical significance (*p* < 0.05).

Beyond these changes in firing dynamics, our analysis revealed a profound effect of the rTMS protocols on the composition of the active motor unit pool. A one-way ANOVA on the average Motor Unit Action Potential (MUAP) amplitude—a proxy for motor unit size—revealed a highly significant difference between the groups, *F*(2,212) = 17.418, *p* < 0.001. Post-hoc tests confirmed that the mean MUAP amplitude was significantly different between all three conditions (M1 vs. Sham, *p* = 0.009; M1 vs. SMA, *p* = 0.015; and SMA vs. Sham, *p* < 0.001) (Figure 4C). This critical finding indicates that the interventions fundamentally altered the recruitment strategy, changing which motor units were selected to perform the task.

### M1 and SMA Inhibition Alter Firing Rate Patterns

We next characterized the dynamic evolution of motor unit firing rate (MUFR) throughout the NFB phase by performing a two-way repeated-measures ANOVA, with Time (six 5-second bins) as the within-subjects factor and Group (M1, SMA, Sham) as the between-subjects factor. A Greenhouse-Geisser correction was applied due to a violation of the sphericity assumption (*χ²*(14) = 11,750.05, *p* < 0.001). The analysis revealed a highly significant Time x Group interaction, *F*(4.78, 5274) = 55.59, *p* < 0.001. This finding indicates that the rTMS protocols dynamically altered the pattern of neural drive over time. To deconstruct this interaction, we conducted planned post-hoc tests comparing each stimulation condition to Sham. As illustrated in Figure 5, these tests confirmed that both M1 and SMA stimulation led to a sustained reduction in MUFR, but with subtly different temporal profiles. For the M1 group, rTMS resulted in a robust reduction in MUFR that was sustained through the middle of the NFB phase, with significant differences observed at the 5-10s, 10-15s, and 15-20s intervals (all *p* < 0.001; Figures 5A and 5B). In comparison, the reduction following SMA inhibition was similarly pronounced but appeared to emerge earlier and persist longer. As shown in Figures 5C and 5D, the SMA group exhibited significantly lower firing rates than the Sham group across a broad window spanning the 1-25s period (all p < 0.03). Collectively, these results demonstrate that while both M1 and SMA inhibition lead to a significant reduction in the magnitude of the neural drive, their distinct temporal profiles provide further evidence for their different roles in shaping the motor command over time.

**Figure 5.**
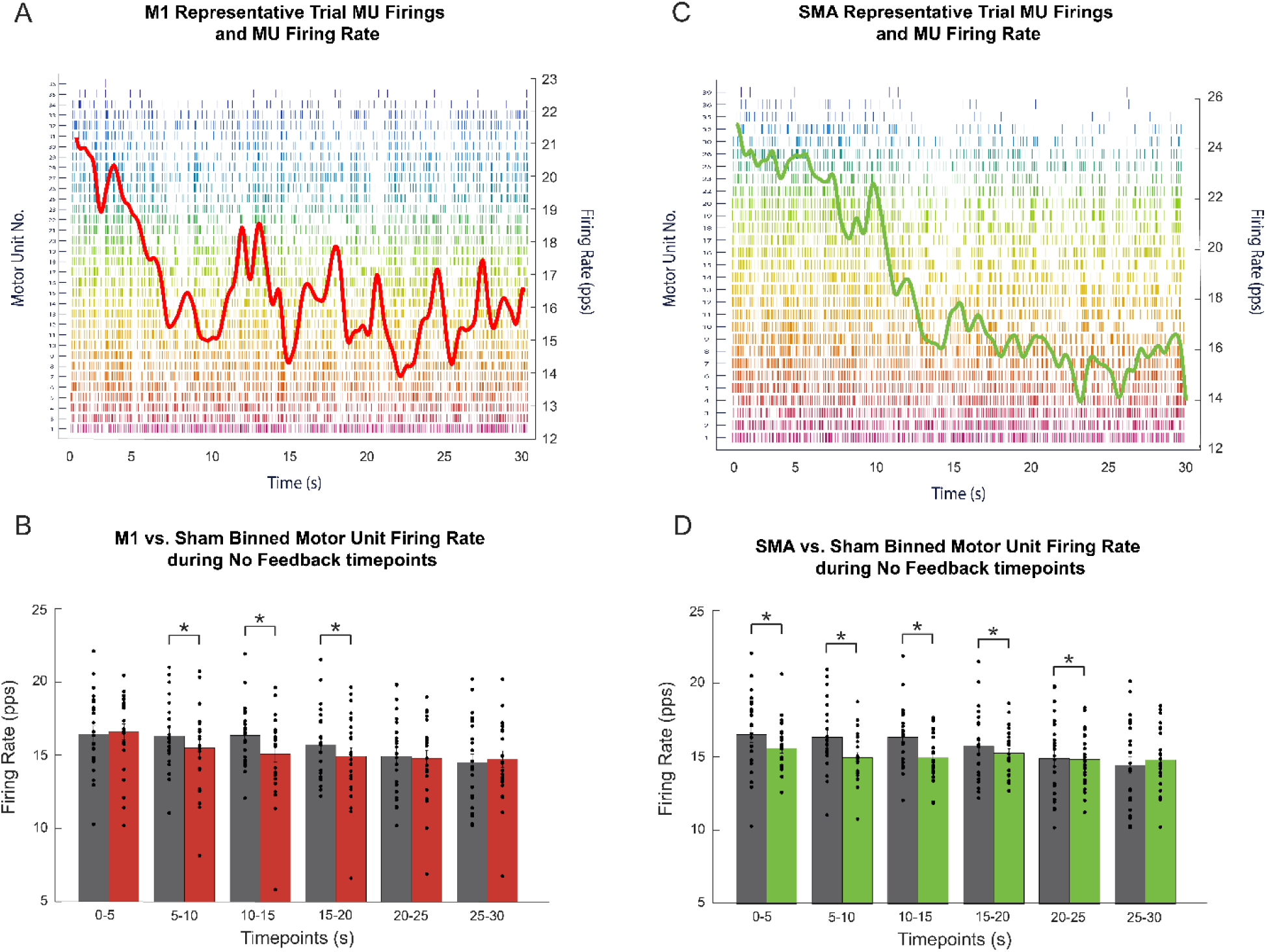
Dynamic Effects of M1 and SMA Inhibition on Motor Unit Firing Rate. This figure illustrates the time course of the neural drive during the NFB phase, revealing distinct temporal profiles for the effects of M1 and SMA inhibition. Representative raster plots from single trials for a participant in the M1 group (**A**) and SMA group (**C**). Each row represents a single motor unit, and each vertical tick marks a firing event. The overlaid continuous line (red/green) shows the smoothed average firing rate across the entire population. Group-averaged binned MUFR for the M1 vs. Sham groups (**B**) and the SMA vs. Sham groups (**D**). Post-hoc tests revealed that both M1 and SMA stimulation significantly reduced MUFR compared to Sham, but with different temporal patterns. Error bars represent SEM, and asterisks (*) denote a significant difference (*p* < 0.05) between the stimulation group and Sham within that time bin.

### M1 Inhibition Creates a Tighter Coupling

To directly test the extent to which the observed behavioral outcomes were a direct consequence of the changes in neural drive, we performed a Pearson correlation analysis between the MUFR and the mean force (%MVC) during the NFB phase for each experimental group. As depicted in Figure 6, a significant positive correlation was found for all three groups. The M1 group exhibited a remarkably strong, near-perfect linear relationship (*r* = 0.96, *p* < 0.001). This indicates that the variance in force output was almost entirely accounted for by the modulation of motor unit firing rate (Figure 6A). This tight coupling provides strong evidence that following M1 inhibition, the dominant mechanism determining force output is the overall magnitude of the neural drive. In contrast, this coupling was substantially weaker in both the SMA group (*r* = 0.82, *p* < 0.001) and the Sham group (*r* = 0.75, *p* < 0.001) (Figures 6B and 6C). This weaker correlation is a critical finding, as it suggests that in these conditions, force output was influenced by a more complex combination of factors beyond just the mean firing rate.

**Figure 6.**
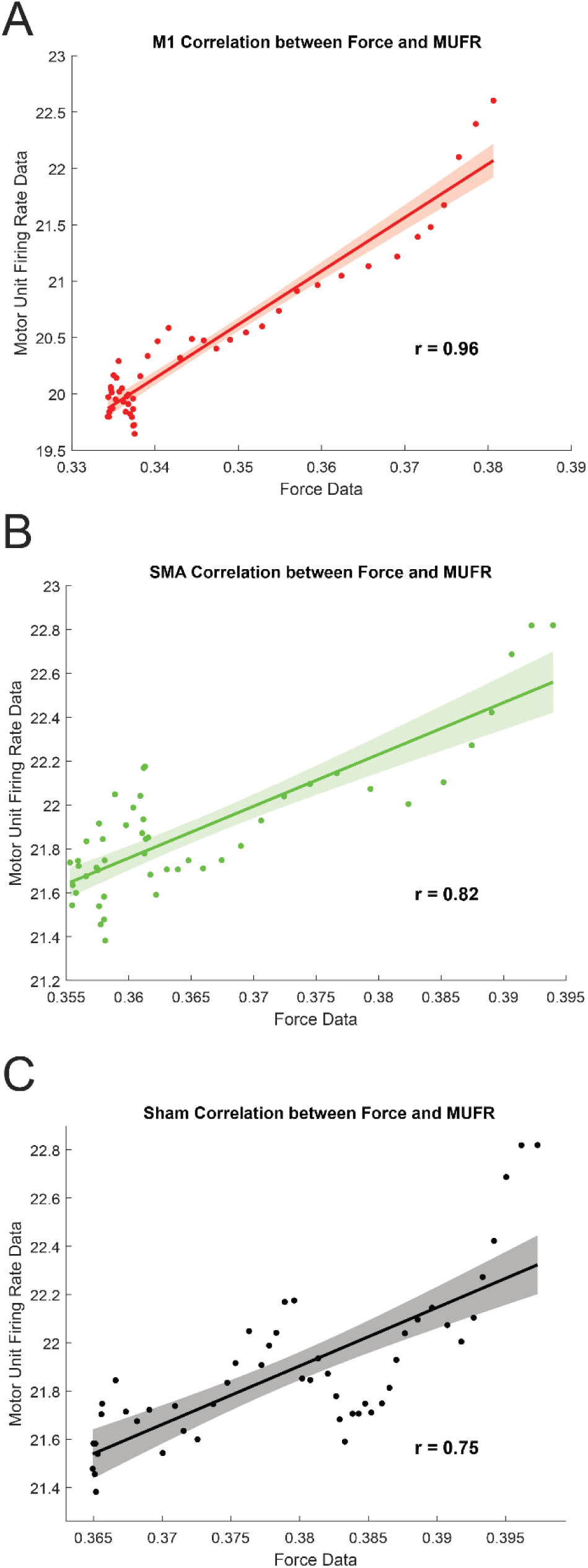
Relationship Between Force and Motor Unit Firing Rate. This figure directly links the behavioral output (force) to the neural drive (firing rate) for each experimental group during the NFB phase. Each panel displays a scatter plot where each dot represents an individual participant’s mean %MVC and mean MUFR. The solid line represents the linear regression fit, and the shaded area is the 95% confidence interval. The Pearson correlation coefficient (r) is displayed for each group. (**A**) The M1 group exhibited a near-perfect linear relationship (r=0.96), indicating a tight coupling between force and firing rate. (**B, C**) This coupling was substantially weaker in the SMA group (r=0.82) and the Sham group (r=0.75), suggesting that other factors, such as recruitment and temporal organization, were also contributing to force production in these conditions.

### SMA Inhibition Uniquely Degrades Coherence

Finally, we analyzed motor unit coherence, a measure of the shared, oscillatory input to the motoneuron pool. A one-way ANOVA revealed a significant effect of group on motor unit coherence, but this effect was specific to the low-frequency Delta band (0-5Hz), *F*(2, 71) = 4.123, *p* = 0.020. In contrast, no significant group differences were found in the Alpha (5-13Hz) or Beta (13-30Hz) bands, indicating that the effect of SMA inhibition was not a general disruption of all oscillatory activity but was targeted to a specific organizing signal. As shown in Figure 7, post-hoc tests to deconstruct the significant Delta-band effect revealed that it was driven by the SMA group having significantly lower coherence than the M1 and Sham group (*p* < 0.03). This finding is critical, as it provides direct neurophysiological evidence that inhibitory rTMS over the SMA uniquely disrupts the organized, low-frequency common drive. This rhythmic signal is thought to be essential for coordinating the firing of the motor unit pool to produce a stable and steady force output during isometric contractions.

**Figure 7.**
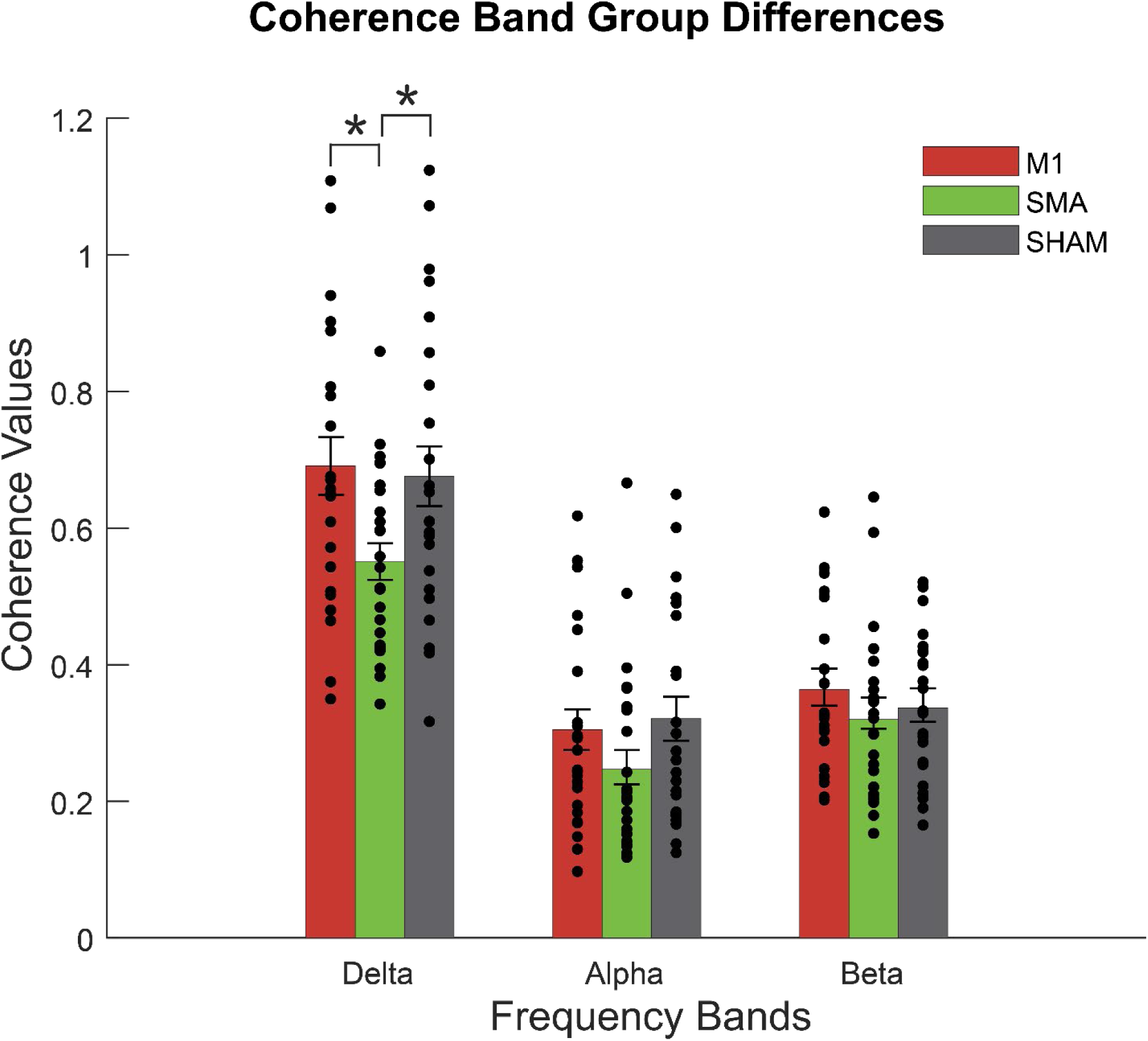
SMA Inhibition Uniquely Degrades Low-Frequency Motor Unit Coherence. This figure presents the effect of the rTMS interventions on the temporal organization of the motor command, as measured by motor unit coherence. The bar graph shows the mean coherence values for the M1 (red), SMA (green), and Sham (gray) groups across three canonical frequency bands: Delta (0-5 Hz), Alpha (5-13 Hz), and Beta (13-30 Hz). A significant group difference was found only in the Delta band. Post-hoc tests revealed that the SMA group exhibited significantly lower Delta-band coherence than the M1 group. This finding indicates that SMA inhibition uniquely disrupts the organized, low-frequency common drive thought to be essential for stabilizing force output. Bars represent the group mean, error bars represent SEM, and asterisks (*) denote a statistically significant difference (*p* < 0.05) between groups.

## Discussion

This study employed inhibitory rTMS to causally dissociate the roles of the primary motor cortex (M1) and supplementary motor area (SMA) in sustained isometric force control at the single motor unit level. Our findings reveal a clear functional divergence. M1 inhibition produced a direct failure of motor output, manifesting as a rapid force decline that was almost entirely explained by a tightly coupled reduction in motor unit firing rates. In contrast, SMA inhibition did not impair the net magnitude of force but instead compelled a complex reorganization of the underlying neural control strategy. This was characterized by a reliance on smaller-amplitude motor units and a marked degradation of the low-frequency (delta-band) coherence that organizes stable motor commands. These results provide direct causal evidence for a functional division of labor where M1 directly governs the magnitude of the motor command, while SMA orchestrates its composition and temporal structure.

Our initial finding—that the removal of visual feedback universally led to a decline in mean force and an increase in force variability across all three experimental groups— aligns with a well-documented phenomenon in motor control known as unintentional force drift (Slifkin et al., 2000; Vaillancourt and Russell, 2002). While this drift has been traditionally attributed to factors like motor memory decay or fatigue, Ambike et al. (2016) provided a more mechanistic explanation rooted in the equilibrium-point hypothesis. According to this framework, the central nervous system does not directly control force, but rather specifies a hypothetical spatial "referent coordinate" for the effector. The isometric force we measure is a consequence of the physical constraints that prevent the finger from reaching this centrally-commanded position. Ambike and colleagues demonstrated that when visual feedback is removed, this referent coordinate begins to slowly drift back toward the finger’s actual physical location, a process they describe as a natural relaxation of the system toward a lower energy state. This drift in the underlying control variable, not just a failure of memory, manifests as the slow decline in force output that we observed. Therefore, the degraded performance seen in all groups during the No Visual Feedback phase represents the baseline challenge of maintaining a stable internal motor command against this natural drift. The distinct effects of M1 and SMA stimulation can thus be interpreted as differential disruptions to the cortical mechanisms responsible for generating and stabilizing this internal command.

The profound impact of M1 inhibition on force production provides direct causal support for its canonical role as the primary and most direct cortical output station for voluntary motor commands. Anatomically, M1 is uniquely positioned for this function, containing a high proportion of corticospinal neurons that project monosynaptically to spinal motor neurons, affording it powerful and direct control over muscle activation (Lemon, 2008). Our finding that disrupting M1 activity impairs force maintenance aligns with numerous studies that have used brain stimulation techniques to demonstrate M1’s critical role in motor execution (Chen et al., 1998; Ziemann et al., 1996; Rizvi et al., 2023; Kallioniemi et al., 2025). While extensive prior work has established a strong *correlation* between M1 activity and force output, from single-unit recordings in primates to human neuroimaging (Evarts, 1968; Omrani et al., 2017), our study extends this understanding by providing direct *causal* evidence in humans at the motor unit level. This causal link is powerfully underscored by the strong correlation (*r* = 0.96) between the decline in force and the reduction in motor unit firing rates. This tight coupling demonstrates that M1’s principal role in this context is to determine the overall magnitude, or gain, of the descending neural drive. When this primary drive is compromised, the motor system appears unable to compensate, resulting in an immediate and unavoidable behavioral failure.

In contrast to the direct motor failure seen with M1 inhibition, the effects of SMA stimulation support its proposed role as a higher-order motor area, critical for motor planning and the implementation of internal models (Roland et al., 1980; Nachev et al., 2008). Neuroimaging studies have consistently implicated the SMA in the preparation and sequencing of actions, particularly when movements must be guided by internal representations rather than external cues (Tanji, 2001). Our central finding—that participants could maintain their target force despite SMA inhibition—provides direct causal support for this framework. This indicates that the fundamental capacity for motor execution via M1 remained intact, but the strategy used to achieve this goal was fundamentally altered. This was evidenced at the motor unit level by a complex reorganization of the neural command. The degradation of delta-band (0-5Hz) coherence is a key finding, as this low-frequency common drive is thought to be essential for organizing and stabilizing motor unit firing during steady contractions (De Luca et al., 1982; Contessa et al., 2016). Its disruption points to a breakdown in the temporal structure of the motor command. Furthermore, the observed reliance on smaller-amplitude motor units suggests that the SMA is involved in selecting the most efficient population of motor units for the task. When the SMA’s plan is disrupted, the motor system appears to compensate by recruiting a different, less optimal pool of motor units. This complex reorganization—altering which units are active and how their firing is timed— explains how force could be maintained even as the underlying command became less efficient. This provides direct causal evidence that the SMA is not a passive relay but is actively responsible for formulating the optimal motor plan that is then implemented by M1.

This functional dissociation between M1 and SMA represents a biomechanically and computationally advantageous control architecture. By segregating the control of command magnitude (M1) from its structural and compositional planning (SMA), the nervous system achieves both robustness and flexibility. M1 can serve as a reliable "gain controller," rapidly adjusting force levels based on direct inputs, while the SMA can formulate and refine motor plans based on task goals and sensory context, such as the absence of visual feedback. The ability of the motor system to compensate for SMA inhibition, albeit inefficiently, causally demonstrates the robustness of the M1 execution module. Conversely, the quality and efficiency of the final motor output are clearly dependent on the integrity of the motor plan provided by the SMA.

Beyond its contribution to fundamental motor neuroscience, this functional dissociation provides a powerful framework for interpreting the pathophysiology of various neurological motor disorders. The direct failure of force magnitude we observed following M1 inhibition closely mirrors the clinical presentation of hemiparesis following a stroke affecting the primary motor cortex. The profound weakness in these patients can thus be mechanistically understood as a failure of this magnitude-control system, where the corticospinal drive necessary to achieve adequate motor unit firing rates is compromised (Ward & Cohen, 2004; Stinear et al., 2007). Therapies aimed at restoring strength, such as high-intensity training or specific non-invasive brain stimulation protocols over M1, are therefore directly targeting this compromised "gain controller." In contrast, our findings suggest that SMA dysfunction produces a distinct clinical profile, characterized not by simple weakness, but by disorganized and inefficient motor control. This aligns well with motor deficits seen in conditions like Parkinson’s disease, where the SMA is known to be hypoactive. Patients with Parkinson’s often struggle with the internal generation and sequencing of movements (akinesia), a deficit in motor planning rather than execution (Jahanshahi et al., 1995; Haslinger et al., 2001). Similarly, conditions like dystonia and certain forms of apraxia are characterized by poorly structured motor commands and an inability to select the correct motor programs, reflecting a failure of the SMA’s organizational role (Hallett, 2011). Our motor unit-level results provide a potential micro-level explanation for the disorganized motor output seen in these disorders. This distinction suggests that therapies for these conditions may be more effective if they focus on providing external cues to bypass internal planning mechanisms or use neuromodulation to specifically target the SMA and its associated networks.

While this study provides novel causal insights, several limitations should be considered when interpreting the results. First, this study employed a between-subjects design, which is susceptible to inter-individual variability in anatomy, physiology, and responsiveness to TMS. Although our baseline measures confirmed that the groups were well-matched on key variables, future studies employing a within-subjects crossover design would provide even more robust evidence by allowing each participant to serve as their own control. Second, our stimulation sites were guided by standard EEG landmarks rather than being co-registered to individual structural MRIs. While our previous work has demonstrated the high spatial focality of stimulation at the 4cm SMA site (Kim et al., 2025), utilizing individual neuroanatomy in future studies would enable a more precise anatomical delineation between subregions like the SMA-proper and pre-SMA, potentially revealing even finer-grained functional specializations. Finally, the present study was intentionally constrained to a relatively simple isometric force task to isolate the fundamental control properties of M1 and SMA. The distinct contributions of these areas may be even more pronounced during complex, dynamic, or sequential movements that place greater demands on motor planning. Investigating how these causal roles evolve during more complex behaviors represents a crucial avenue for future research. Despite these limitations, our findings provide robust causal evidence for a functional division of labor between M1 and SMA, contributing to a more mechanistically detailed understanding of the cortical motor hierarchy.

## Data availability statement

The data that support the findings of this study are available from the corresponding author upon reasonable request.

## Author Contributions

Y.R.E., and Y.L. designed research; Y.R.E. performed research; Y.R.E., and Y.L. analyzed data; and Y.R.E., and Y.L. wrote the paper.

